# Pervasive Chromosomal Instability Drives the Karyotypic Evolution of Hypodiploid Tumours

**DOI:** 10.1101/2025.08.08.668983

**Authors:** Elle Loughran, Aoife McLysaght, Máire Ní Leathlobhair

## Abstract

Tumours frequently exhibit extreme levels of aneuploidy. While increases in ploidy are well-characterised, the opposite phenomenon—extensive chromosome loss leading to hypodiploidy— remains underexplored. Here, we analyse over 17,000 cancer genomes from 34 cancer types and perform a pan-cancer analysis of karyotypic evolution in hypodiploid tumours.

We find that hypodiploidy is widespread and associated with a generalised chromosomal instability phenotype, marked by significantly elevated rates of genome doubling, intrachromosomal copy number alterations, chromothripsis, and intra-tumour heterogeneity. These tumours are hypoxic and strongly enriched for *TP53* mutations. However, we also identify a subset of cancers—acute lymphoblastic leukaemia (ALL), kidney chromophobe, and adrenocortical carcinoma—that exhibit stable hypodiploidy, with stereotyped chromosome loss patterns, low chromosomal instability, and distinct evolutionary origins. We exploit this stability to develop a simple method of distinguishing poor-prognosis masked hypodiploid from good-prognosis hyperdiploid ALL using only cytogenetic data, enabling more precise risk stratification.

Finally, we show that hypodiploidy predicts poor prognosis across cancers. Genome doubling does not confer a fitness advantage in low-hypodiploid tumours, nor do these tumours evolve to avoid loss of dosage-sensitive genes. Together, these findings provide the first pan-cancer characterization of hypodiploidy as a widespread and clinically relevant phenomenon often driven by pervasive chromosomal instability, and illustrate the remarkable ability of cancer cells to tolerate and evolve under extreme dosage imbalance.

## INTRODUCTION

Aneuploidy is a hallmark of cancer, observed in 90% of solid tumours and a majority of haematological cancers.^1,2^ The extent of aneuploidy varies significantly, both within and between cancer types: while many tumours possess one or a few recurrent aneuploidies, often affecting chromosomes carrying specific cancer driver genes,^3,4^ others exhibit widespread aneuploidy across much of the genome, resulting in marked deviations from diploidy. Extreme aneuploidy often results from whole genome duplication (WGD), which is the second most common alteration in cancer and is associated with chromosomal instability, tumour progression, and poor prognosis.^5,6,7^

More rarely, tumours may lose large numbers of chromosomes to reach a hypodiploid karyotype. This phenomenon has been best studied in acute lymphoblastic leukaemia (ALL), where near-haploid (*<*30 chromosomes) and low-hypodiploid (30-39 chromosomes) cases form a rare subgroup with distinct mutational profiles,^8^ poor treatment response^9,10^ and very poor prognosis.^11^ Near-haploid or low-hypodiploid subtypes have also been observed in other cancers including kidney chromophobe carcinoma,^12^ chondrosarcoma,^13^ giant cell glioblastoma,^14^ malignant pleural mesothelioma^15^ and oncocytic follicular thyroid carcinoma.^16^ In at least some tumour types, hypodiploidy can arise early in tumour development^17^ and persist throughout metastatic progression.^18^

Hypodiploid tumours frequently undergo WGD,^19, 13^ resulting in hyperdiploid or near-triploid karyotypes with homozygous disomies and heterozygous tetrasomies. It has been proposed that WGD is positively selected in hypodiploid tumours to normalize gene dosage,^11^ and/or buffer the effects of deleterious mutations in regions with hemizygous deletions or loss of heterozygosity.^20^ In ALL, WGD is rare overall but occurs in 64% of near-haploid and 44% of low-hypodiploid cases.^11^ In some patients, genome-doubled and non-doubled subclones can coexist, with the doubled subclone often dominating at diagnosis and the hypodiploid founder clone at relapse.^2^ Because the doubled clone retains the poor prognosis of its hypodiploid precursor but may be misclassified as a good-prognosis high-hyperdiploid case (‘masked hypodiploidy’), WGD presents an important diagnostic challenge in hypodiploid ALL.^21, 22^

Hypodiploid subtypes in cancer types including ALL,^22^ kidney chromophobe cancer^23^ and Hürthle cell carcinoma^18^ exhibit tissue-specific stereotyped aneuploidy patterns, with recurrent losses of specific chromosomes. The biological basis for these patterns is generally unknown, although it has been suggested that chromosome 7 may be constrained against monosomy in thyroid cancer cells due to the cancer-essential imprinted genes it carries,^24^ and that chromosome 21 is consistently retained in hypodiploid ALL due to its leukemogenic effects.^25^ The mechanisms by which these karyotypes arise are also unclear, but could involve successive losses or multipolar mitosis.^2^ Beyond its prognostic significance, an improved understanding of hypodiploidy thus offers insight into the cellular constraints and selective forces that shape tumour karyotypes in the context of extensive chromosome loss.

In this work, we perform a pan-cancer analysis of hypodiploidy in order to understand the prevalence, origins, and evolution of low-hypodiploid and near-haploid tumours, particularly outside ALL. Using data from 17,239 tumours in The Cancer Genome Atlas (TCGA) and the Mitelman Database of Chromosome Aberrations and Gene Fusions in Cancer,^26, 27^ we identify hypodiploid tumours across a wide range of haematological and solid cancers and analyse patterns of chromosome loss with respect to tissue type and chromosome features. We characterise the relationship between hypodiploidy and chromosomal instability within and between cancer types, and identify genomic and environmental factors associated with hypodiploidy. This work distinguishes classes of stable and unstable hypodiploid tumours, maps an underlying chromosomal instability phenotype that connects copy number alterations at multiple scales, and contributes to our knowledge of the origins of extreme aneuploidy.

## RESULTS

### Hypodiploidy is Widespread Across Cancer Types

To understand patterns of hypodiploidy pan-cancer, we began by identifying hypodiploid cases across 33 cancers from TCGA and comparing their chromosome count distributions to those observed in hypodiploid ALL. We first established a reference distribution for hypodiploid chromosome content by analyzing 7,922 clones from 6,907 karyotyped ALL cases in the Mitelman Database. Using CytoConverter, we inferred copy number profiles from karyotype strings and visualised the distribution of autosome counts in ALL cases with fewer than 44 autosomes (Fig. 1a). As well as a large near-diploid peak for cases with rearrangements that create dicentric chromosomes,^2^ we observed a low-hypodiploid peak with 32-37 autosomes (129 clones) and a near-haploid peak with 23-27 autosomes (165 clones). Only 15 clones fell into the 28-31 autosome range between the two peaks. This distribution recapitulates the known bimodal distribution of chromosome counts in hypodiploid ALL.^11^ The lowest autosome count was 23, observed in 18 near-haploid cases with disomy 21.

**Figure 1:**
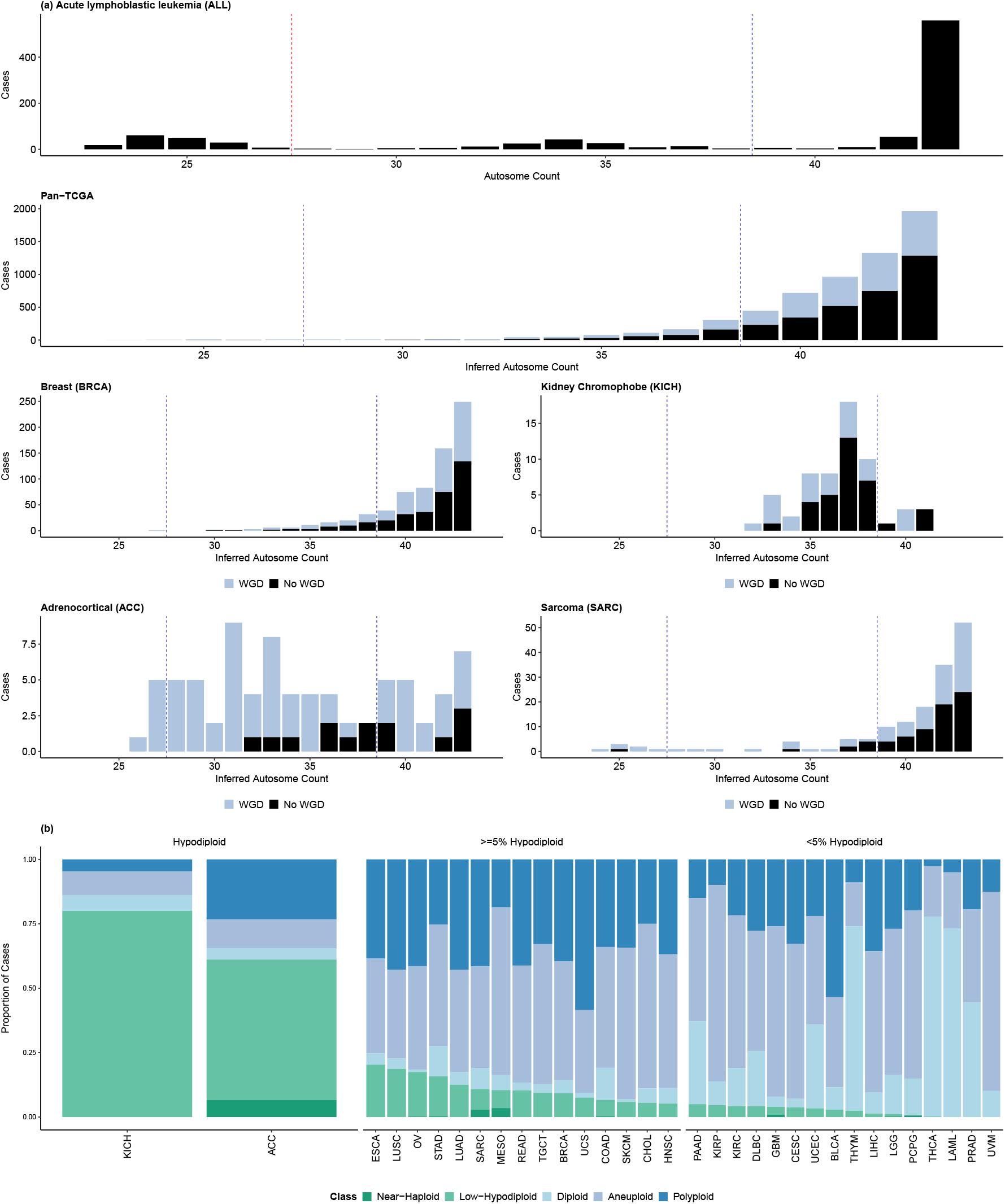
Prevalence of hypodiploidy in the TCGA and Mitelman datasets. **A**, Inferred sub-diploid (*<* 46 chromosomes, *<* 44 autosomes) autosome count distributions for selected cancer types. Minimal autosome counts were inferred in the TCGA dataset based on loss of heterozygosity to identify former hypodiploids which have since gained chromosomes or undergone WGD. The ALL plot is based on cytogenetic data and shows only current hypodiploids. Dashed lines indicate our thresholds for low-hypodiploidy and near-haploidy. **B**, Proportions of each TCGA cancer type that are near-haploid (*<* 28 autosomes), low-hypodiploid ≤ 38 autosomes), diploid (all autosomes disomic), polyploid (positive WGD call and no hypodiploid history), and aneuploid (all other cases). Current and former hypodiploids are both counted as hypodiploid. Tumour abbreviations are reported as per https://gdc.cancer.gov/resources-tcga-users/tcga-code-tables/tcga-study-abbreviations.

We additionally examined the distribution of autosome counts in kidney chromophobe cancer, a TCGA cancer type with an established hypodiploid subtype,^28, 29^ and found that sub-diploid cases clustered around 35-38 autosomes. We used the distributions of autosome counts in ALL and KICH to set the threshold for hypodiploidy in this study at≤ 38 autosomes, which corresponds to ≥ 6 autosome losses. This threshold resulted in the identification of 175 TCGA cases (1.7%) as hypodiploid at time of sequencing. To account for cases where hypodiploidy may have been masked by genome doubling or subsequent chromosome gains, we also identified an additional 646 cases (6.3%) with loss of heterozygosity (LOH) across ≥ 6 autosomes but autosome counts above 38, a genomic signature consistent with past hypodiploidy.

In contrast to the bimodal distribution observed in hypodiploid ALL, autosome losses in the TCGA cohort were exponentially distributed, with karyotype frequency decreasing with the degree of genomic loss. In line with this, low-hypodiploidy was far more common than near-haploidy: of the 821 current and former hypodiploid cases (LOH of ≥ 6 autosomes), 96.7% fell within the low-hypodiploid range. Near-haploid cases were identified across 10/33 cancer types but were extremely rare (27/10,332 TCGA cases, 0.26%), reaching a prevalence ≥ 1% only in ACC (6/90), mesothelioma (3/86) and sarcoma (7/248). Most of these cases (24/27) had undergone genome doubling, resulting in a wide range of autosome counts (26-117) at time of sequencing.

The prevalence and distribution of hypodiploid karyotypes varied widely across cancer types (Fig. 1b). Low-hypodiploidy was most common in KICH and adrenocortical carcinomas (ACC), at 80% (52/65 cases) and 54.4% (49/90 cases), respectively. Low-hypodiploidy was rarer in other cancer types but widespread, with 24/33 TCGA cancer types having at least five low-hypodiploid cases and esophageal, lung squamous, ovarian, stomach, lung adenocarcinoma and rectal adenocarcinoma being ≥ 10% low-hypodiploid. Several cancer types other than KICH diverged from the pan-cancer exponential trend, with sarcomas showing a notable surplus of near-haploid cases and ACC presenting a distinctly uniform distribution of mostly genome-doubled low-hypodiploids and near-haploids. Two blood cancers are represented in the TCGA dataset; no hypodiploids were identified among the acute myeloid leukaemia cases, and hypodiploidy was uncommon in the diffuse B-cell lymphomas (2/47 cases, 4%).

### MH Score Identifies Masked Hypodiploidy in ALL with High Specificity

Genome doubling (WGD) is also common in hypodiploid ALL, resulting in “masked hypodiploids” with hyperdiploid or near-triploid karyotypes resembling high hyperdiploid ALL (HeH-ALL). These cases present a significant diagnostic challenge. Masked hypodiploid acute lymphoblastic leukemia (MH-ALL) retains a very poor prognosis but may be misclassified as HeH-ALL, a subtype with good prognosis and favourable treatment response, if the underlying hypodiploidy remains undetected.^11^ Distinguishing these two divergent subsets using only the cytogenetic data available in routine clinical investigations is thus of both biological and clinical significance.

Masked hypodiploids show a characteristic pattern of ‘hyperdiploidy by tetrasomy’, with their ‘hyperdiploid’ karyotype being made up of disomies and tetrasomies (Fig. 2a) rather than the disomies and trisomies seen in HeH-ALL.^22^ We hypothesized that masked hypodiploid and HeH-ALL could be systematically distinguished by the simple heuristic # *tetrasomies - # trisomies*, where a masked hypodiploidy (MH) score *>* 0 indicates MH-ALL and a score *<*= 0 HeH.

**Figure 2:**
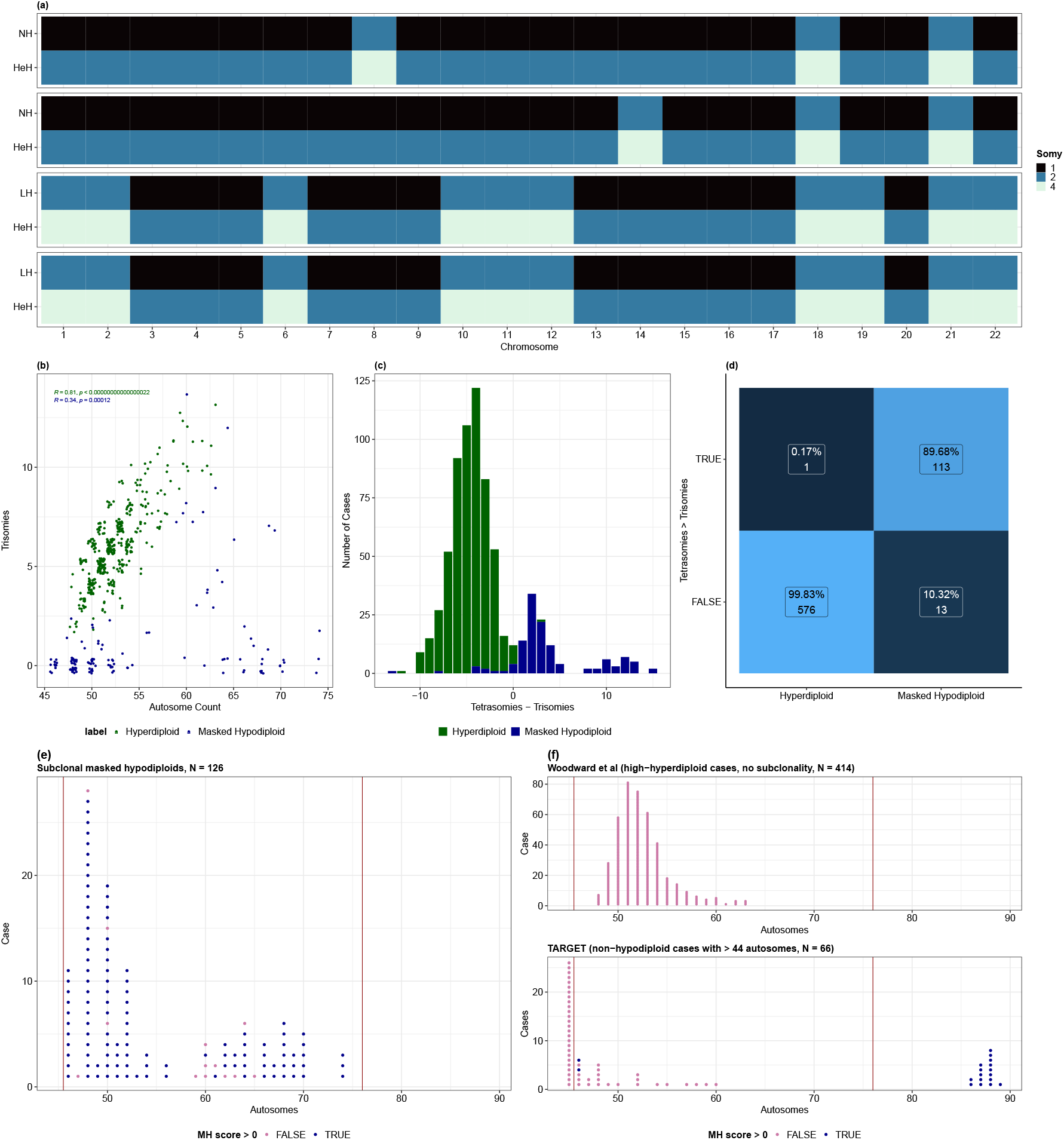
Identification of masked hypodiploid ALL based on cytogenetic data. **A**, Visualisation of genome doubling leading to subclonal masked hypodiploidy in two low-hypodiploid and two near-haploid ALL cases from the Mitelman database. Patients represented include a two-year-old female, a fifteen-year-old male, a forty two-year old female and a thirteen-year-old female karyotyped by Harrison et al (2004). NH, near-haploid; LH, low-hypodiploid; HeH, hyperdiploid. **B**, Relationship between total autosome count and number of trisomies in hyperdiploid vs masked hypodiploid ALL. Hyperdiploid cases with subclonal whole chromosomes were excluded. **C**, Distribution of MH score (# tetrasomies - # trisomies) in hyperdiploid vs masked hypodiploid ALL. **D**, Performance of MH score in distinguishing between high-hyperdiploid and masked hypodiploid ALL. **E**, Distribution of autosome counts and MH score classification in subclonal masked hypodiploids from the Mitelman database. **F**, Distribution of autosome counts and MH score classification in non-hypodiploid ALL cases: top, high-hyperdiploid cases from Woodward et al (2023), excluding cases with subclonal variation in chromosome counts; bottom, cases from the TARGET-ALL-P2 project with *>* 44 autosomes. Vertical lines indicate the range of potential doubled hypodiploid autosome counts.

To test this heuristic, we used MEDICC2 to construct copy number phylogenies for 150 multi-clone samples from the Mitelman database and identified 126 subclonal masked hypodiploids. For comparison, we obtained allele-specific copy number data for 577 HeH-ALL cases from Woodward et al (2023)^30^ and examined the distribution of MH scores between the two groups. Trisomies were much rarer in masked hypodiploids: the median MH-ALL case had zero trisomies, versus a median of six trisomies in HeH-ALL. There was a strong correlation between the total number of autosomes and the number of trisomies in HeH-ALL (r = 0.81) but not MH-ALL (r = 0.34) (Fig. 2b). MH scores for the two groups showed little overlap (Fig. 2c), and the heuristic identified masked hypodiploids with a specificity of 99.8% and a sensitivity of 89.7% (Fig. 2d).

This test is simple and applies to both near-haploid and low-hypodiploid ALL. However, it is specific to the hyperdiploid/near-triploid range and will not detect hypodiploids that have undergone two WGDs, which could be problematic if the likelihood of a first WGD and of a second are not independent. Using allele-specific data for 293 ALL cases from the TARGET-ALL-P2 project, we could not find any examples of this scenario: 0/16 cases in the near-tetraploid range had widespread LOH suggestive of a hypodiploid history (Fig. S1a). The sensitivity of the MH score heuristic additionally depends on genome stability post-WGD, as gains from disomy and losses from tetrasomy reduce signal. Across the TCGA dataset, which is dominated by chromosomally-unstable tumours, there is a significant overlap between the MH scores of doubled hypodiploids and those of hyperdiploids, reducing discriminative power (Fig. S1b). The 13 false negative cases in our analysis (non-hypodiploid and hypodiploid karyotypes in the same sample but MH score ≤ 0) had a median MH score of −3 and 7 trisomies (Fig. S1c). Four had an MH score of exactly zero. Some of these samples may have been misassigned to the masked hypodiploid test set, i.e. classified as masked hypodiploids when the coexisting hypodiploid and non-hypodiploid populations in the sample were not related by a genome doubling event. However, they may also represent a subtype of MH-ALL that is chromosomally unstable after genome doubling. While low-hypodiploids form a minority of the test masked hypodiploids (45/126, 35.7%), they make up 61.5% (8/13) of the false negative cases, suggesting that low-hypodiploids may be more chromosomally unstable than near-haploids. Most of the false negatives were in cases with chromosome counts in the good-prognosis high-hyperdiploid range, causing sensitivity in this range to fall to 82.5% (Fig. 2e).

### MH Score Detects Hidden Hypodiploid Origins in Hyperdiploid ALL

We used the MH score heuristic to search the Mitelman database for fully-masked hypodiploid ALL cases, i.e. cases where the original hypodiploid clone has been lost and only the genome-doubled clone remains. We identified 13 fully-masked cases in the high-hyperdiploid range (49-65 autosomes); all were clonal except for one case with both a 50-autosome and a 47-autosome subclone. Another 69 hyperdiploid cases possessed a hypodiploid subclone, suggesting that approximately 8.3% of the 989 hyperdiploid ALLs in the Mitelman database may have hypodiploid origins, including 1.3% which are fully masked. This is likely to be an underestimate, given the high specificity (99.8%) and moderate sensitivity (82.5%) of the MH score in the high-hyperdiploid range.

Broadening our search across the full range of potential doubled hypodiploid autosome counts (46-76), we identified an additional 48 cases (50 clones) with MH score *>* 0. Most (46/48) were within the 46–65 autosome range, with only two cases falling in the 66–76 range. This supports the hypothesis that less-extreme hypodiploids rarely undergo WGD or, when they do, the genome-doubled clone seldom becomes fixed within the tumour population. The vast majority of fully-masked cases had autosome counts of 46 (42 clones) or 48 (6 clones) (Fig. S1d). This differs substantially from the distribution of autosome counts in non-doubled hypodiploid ALL (Fig. 1a) and subclonal genome-doubled hypodiploid ALL (Fig. 2e). In the TARGET-ALL-P2 cohort, only 4/6 (66%) 46-autosome cases were correctly assigned as non-hypodiploid (Fig. 2f). While data in the low-hyperdiploid range is limited, these findings suggest that the specificity of the MH score may decrease at the lower end of the range of potential doubled hypodiploid autosome counts.

### Chromosome Loss Patterns in Hypodiploid Tumours

Having identified hundreds of current and former hypodiploid tumours across 31 cancer types, we next investigated patterns of chromosome loss in hypodiploid ALL and the TCGA tumours. Focusing on low-hypodiploid tumours, we found that whole chromosome and chromosome arm loss rates were non-uniform in all 16 cancer types with at least 15 low-hypodiploids (chi-square test of uniformity, *p <* 1e-4 for each cancer type), with the strongest deviations from uniformity seen in ALL, KICH and ovarian serous cystadenocarcinoma (OV). While we observed some cross-tissue patterns, including frequent retention of chromosomes 7, 1 and 20 and recurrent loss of chromosomes 13 and 17, significant inter-tissue variation remained (Fig. 3a). To investigate the extent to which these non-random chromosome loss and retention patterns in hypodiploid tumours are tissue-specific, we clustered low-hypodiploid samples by their chromosome arm loss profiles. Three cancer types stood out for forming clusters and having higher intra-tissue similarities than their highest inter-tissue similarity: KICH, ALL, and ACC (Fig. 3b). OV also formed a visible cluster but had slightly more similarity to UCEC than to other OV cases. Other cancer types did not form clear clusters and showed higher median similarity with at least one other tissue than with their own samples (Fig. S3c). This suggests that, in low-hypodiploids from most cancer types, chromosome loss is largely stochastic and/or driven by factors common across multiple cancer types rather than tissue-specific selection.

**Figure 3.**
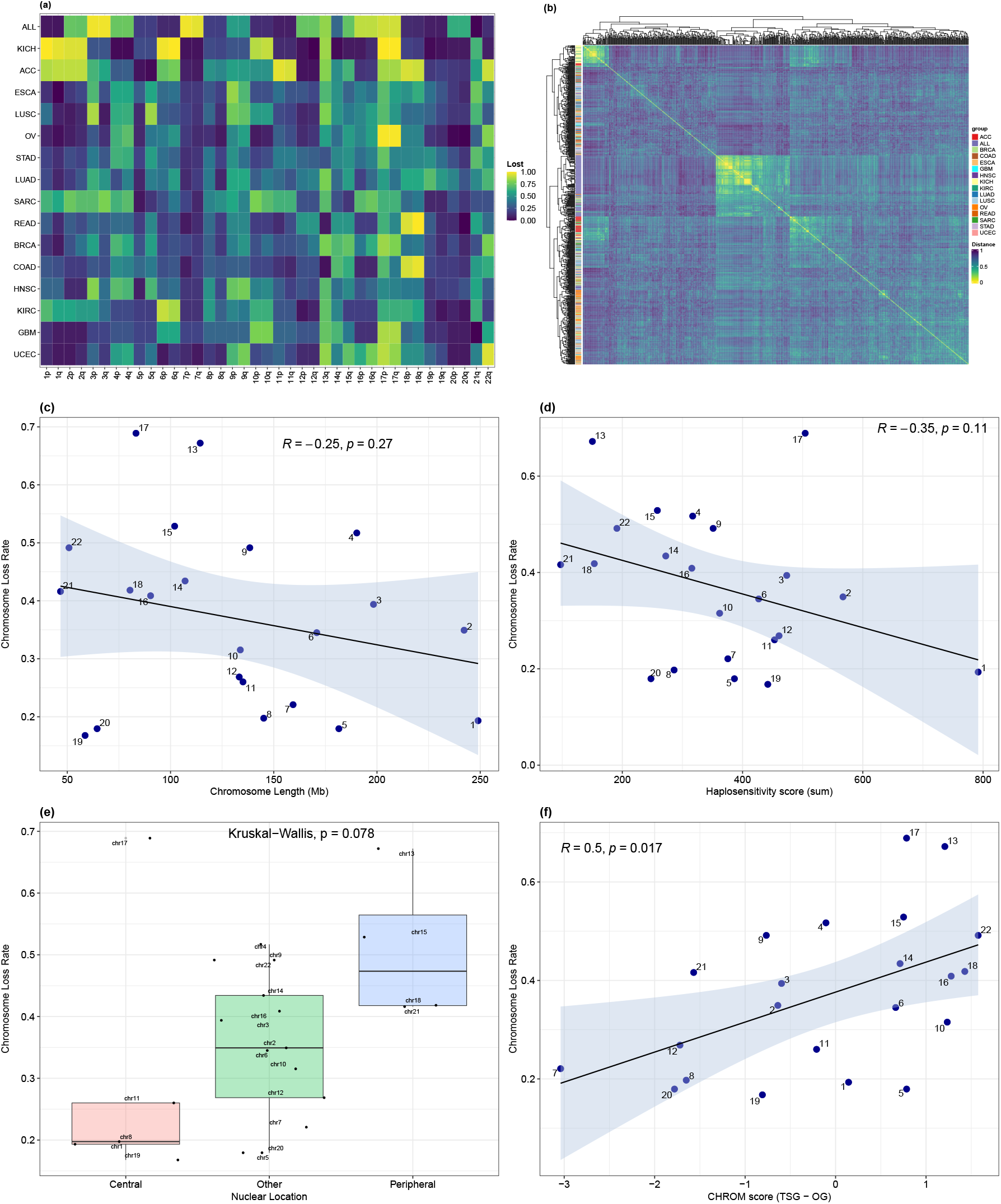
Patterns of chromosome loss in hypodiploid tumours. A, Chromosome arm loss rates among lowhypodiploids in cancer types with ≥ 15 low-hypodiploid cases. B, Clustering of hypodiploid tumours based on chromosome arm loss patterns. Distance was computed using the Jaccard index. Correlation between pan-cancer chromosome loss rates and C, chromosome length, D, sum of gene haplosensitivity probabilities derived from Collins et al (2022), E, chromosome nuclear position as defined by Pari et al (2023) based on translocation frequencies, points jittered horizontally for visibility F, weighted CHROM (TSG - OG) score from Davoli et al.

Whole-chromosome losses predominated over arm-level events in low-hypodiploids, with strong correlations between loss rates for arms of the same chromosome (r = 0.78, *p* = 0.00024). Inter-arm loss rate correlations in less extreme aneuploid samples were significantly weaker, falling to r = 0.27 in tumours with one chromosome loss (Fig. S2b). Even among hypodiploid TCGA cases, however, there were outlying chromosomes with diverging loss rates between arms: 5q was lost in 46% of hypodiploid cases while 5p was only lost in 19%, and chromosome 3 showed the opposite pattern, with the p-arm being lost in 53.2% of hypodiploid cases and the qarm being lost in 31.4%. These patterns suggest selection to retain specific arms despite the overall genomic bias towards chromosomal loss.

### Driver Gene Density Is Dominant Predictor of Chromosome Loss

To explain non-random chromosome loss patterns in low-hypodiploids, we evaluated chromosome features including length, nuclear location, driver gene density, and dosage sensitivity. Across cancer types, we observed no significant correlation between loss rate and chromosome length (Fig. 3c) or dosage sensitivity scores (Fig. 3d). It has been suggested that chromosome 7 is almost always retained in duplicate even in near-haploid cancers because of selection to maintain heterozygosity of cancer-essential imprinted genes;^24^ 7 was indeed the most-retained chromosome among the low-hypodiploid and near-haploid TCGA tumours, being lost in only 9.74% of cases, but was lost in 95.9% of low-hypodiploid and 100% of near-haploid ALL cases.

We next considered the role of chromosome nuclear positioning, using classifications based on translocation frequencies from Parl et al. (2023).^31^ Chromosomes located towards the periphery of the nucleus showed a non-significant trend toward higher loss rates: in low-hypodiploids, these chromosomes had a median pan-cancer loss rate of 47.3%, compared to 19.7% for centrally-located chromosomes and 34.9% for chromosomes with intermediate positioning (Fig. 3e). This trend became significant when we removed chromosome 17, an outlier which carries *TP53* and *BRCA1* and has a much higher loss rate than other centrally-located chromosomes (Fig. S2d).

Of all features tested, driver gene density was the most consistent and significant predictor of chromosome loss. Chromosome-wide TSG-OG density scores derived from Davoli et al. (2013)^32^ had a significant positive correlation with chromosome loss rate in hypodiploid tumours (r = 0.5, *p* = 0.017, Fig. 3f). In a tissue-specific multivariate regression considering all features, driver gene density was the only significant predictor of chromosome loss rate (Fig. S2e). However, within most individual cancer types, none of the tested features, including driver density, significantly predicted chromo-some loss rates, and a pan-cancer multivariate regression showed no significant association between loss rate and any feature (driver gene density *p* = 0.053). This pan-cancer result appears to be confounded by the predominance of ALL cases among low-hypodiploids; when ALL is excluded, the correlation between driver gene density and loss rate strengthens (r = 0.62), and the association is statistically significant across the remaining tissues (*p* = 0.0059).

To determine whether features associated with chromosome loss varied with the degree of aneuploidy, we stratified aneuploid TCGA samples based on the number of chromosomes with LOH (from 1 to 6+, indicating hypodiploid tumours). Haplosensitivity score and peripheral location were both individually associated with chromosome loss rate in less extreme aneuploid samples with 1-5 chromo-some losses (negatively and positively, respectively). However, in multivariate analyses, the TSG - OG density score was consistently the only significant predictor of loss rate pan-cancer (Fig. S2f).

### Genome Doubling Increases with Degree of Hypodiploidy

Low-hypodiploid tumours are highly aneuploid. Although aneuploidy is strongly associated with chromosomal instability (CIN), the inability to maintain a consistent chromosome complement across cell divisions, some highly aneuploid cancer sub-types such as high-hyperdiploid ALL are chromosomally stable.^30^ We thus sought to quantify and characterise the extent of different manifestations of CIN in hypodiploid tumours, including failures of ploidy maintenance (WGD), small-scale intrachromosomal copy number alterations, structural alterations (chromothripsis) and intra-tumour copy number heterogeneity.

Using the MH score heuristic, we first estimated the rate of genome doubling in hypodiploid versus non-hypodiploid ALL in the Mitelman database. WGD was most frequent in near-haploid tumours (67.6%), followed by low-hypodiploids (37%), and almost absent in non-hypodiploid tumours (1.19%) (Fig. 4a). When MH scores for clones with autosome counts outside the well-characterised high-hyperdiploid range were discounted (see Methods), the proportions decreased to 58.8% of near-haploids, 36.1% of low-hypodiploids, and 1.21% of other tumours, but the pattern was preserved (Fig. S3a).

**Figure 4:**
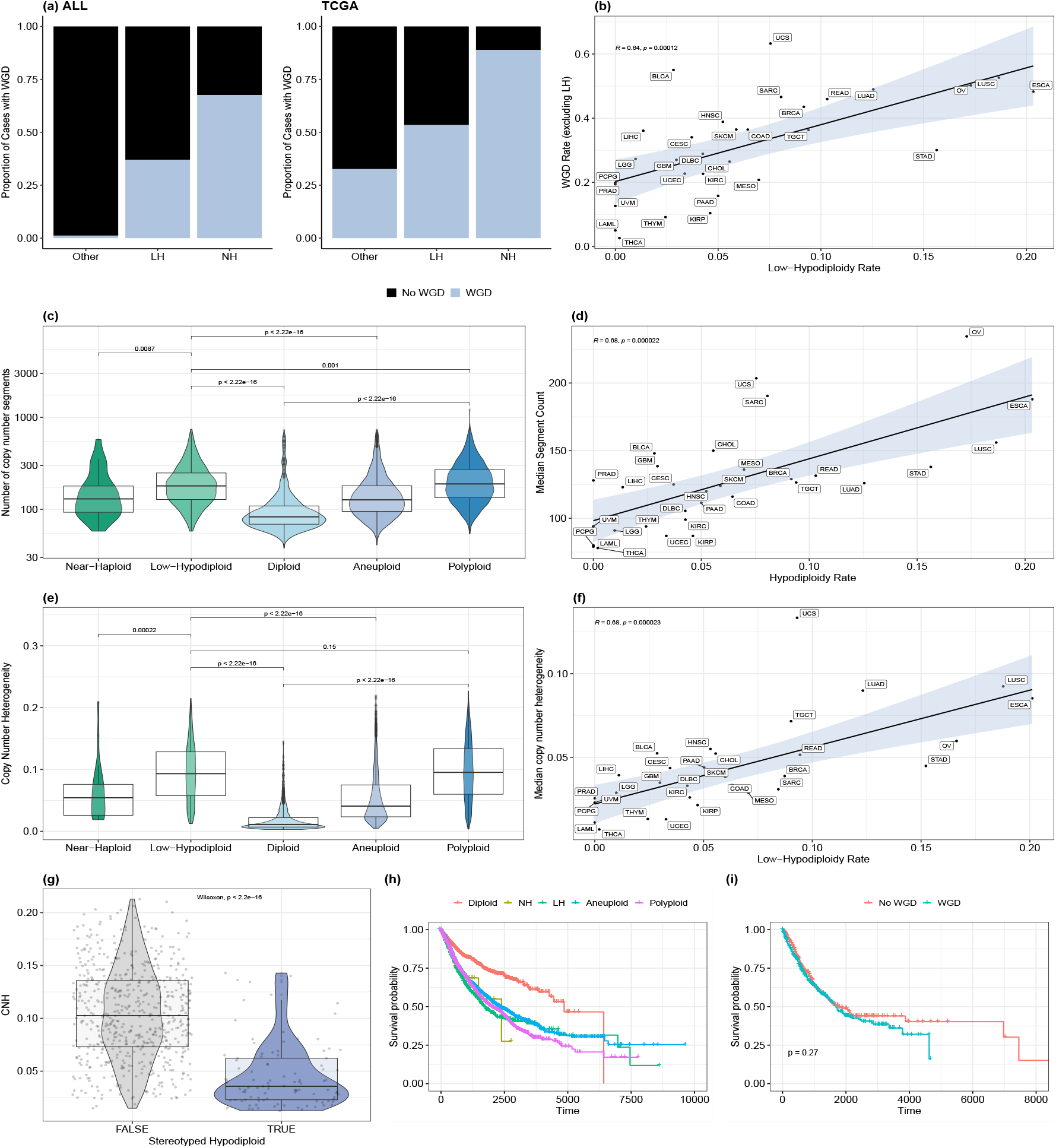
Hypodiploid tumours are distinguished by chromosomal instability at multiple scales. **A**, WGD rate by hypodiploidy level in ALL and the TCGA dataset. Multi-clone cases in ALL were assigned their lowest ploidy class. **B**, Correlation between rate of low-hypodiploidy and WGD rate in non-low-hypodiploid tumours across TCGA cancer types. **C**, Distribution of log10-transformed number of copy number segments by ploidy class. WGD-high indicates cases with a WGD event and no hypodiploid history. **D**, Correlation across TCGA cancer types between rate of low-hypodiploidy and median number of segments per cancer type. Median number of segments was calculated across non-ploidy-altered (diploid and aneuploid) cases. **E**, Distribution of copy number heterogeneity (CNH, van Dijk et all) across ploidy classes. **F**, Correlation between rate of low-hypodiploidy and median CNH across TCGA cancer types. Median CNH was calculated across non-ploidy-altered cases. **G**, Distribution of CNH in low-hypodiploid cases from cancer types with stereotyped and non-stereotyped chromosome loss patterns (KICH and ACC vs BRCA, LUSC, OV, STAD, LUAD, ESCA, COAD, HNSC, KIRC, SARC, UCEC, READ, GBM). **H**, Overall survival by ploidy class. NH, near-haploid; LH, low-hypodiploid. **I**, Overall survival in patients with doubled vs non-doubled low-hypodiploid tumours. P-value calculated using the log-rank test in ggsurvplot() is not adjusted for covariates; adjusted p-value is reported in text. Figs. 4b-i are based on the TCGA dataset. KICH and ACC were excluded from (b), (d) and (f) due to their extremely high low-hypodiploidy rates.

We observed a similar trend of increasing WGD rate with degree of hypodiploidy in TCGA, with WGD occurring in 88.9% of near-haploids (24/27), 53.4% of low-hypodiploids (424/794), and 32.6% of non-hypodiploid tumours (Fig. 4a). It has been suggested that the link between loss of heterozygosity and WGD is due to selection to buffer LOH of essential regions.^20^ However, this may also be explained by a generalised CIN phenotype underlying both aneuploidy/loss of heterozygosity and genome doubling. Supporting this interpretation, the prevalence of low-hypodiploidy and genome doubling was strongly correlated across cancer types (r = 0.64, *p* = 0.00012), even after excluding low-hypodiploid cases from the WGD rate calculation (Fig. 4b).

### Extensive Intrachromosomal Instability and Heterogeneity in Low-Hypodiploid Tumours

We next considered smaller-scale CIN leading to intrachromosomal copy number alterations (CNAs). Hypodiploid tumours have significantly higher numbers of copy number segments across the genome than diploid tumours (Fig. 4c), suggesting a high rate of CNAs beyond the whole-chromosome aneuploidies by which they are defined. An alternative measure of intrachromosomal copy number alterations, the proportion of a chromosome taken up by its longest contiguous copy number segment, shows a similar result, being lower in hypodiploid and polyploid tumours compared to diploids, indicative of more fragmented genomes (Fig. S3b). These differences remain significant when genome-doubled hypodiploids are excluded from the analysis (Fig. S3c-d).

While hypodiploidy or polyploidy themselves may result in increased intrachromosomal instability, hypodiploidy rate is correlated with median segment count across cancer types (r = 0.68, Fig. 4d), further supporting an underlying generalised CIN phenotype. Hypodiploids are also enriched for structural instability, with 55.2% of low-hypodiploids showing evidence of chromothripsis, compared to 8.7% of diploids and 35.8% of less-extreme aneuploids, and second only to polyploid tumours (56.5%) (Fig. S3e).

To more directly index CIN, we used a copy number heterogeneity (CNH) metric developed by van Dijk et al (2019),^7^ which proxies intra-tumour heterogeneity in copy number profiles based on distance from integer copy number values in bulk tumour samples. Copy number heterogeneity was significantly increased in low-hypodiploid compared to diploid and less-extreme aneuploid tumours, suggesting ongoing karyotypic evolution and diversity in karyotypes between cells of the same tumour (Fig. 4e). This held even when genome-doubled cases were excluded (Fig. S3f). In line with the strong correlations between hypodiploidy rate and rates of WGD and intrachromosomal CNA, hypodiploidy rate was strongly correlated with median CNH across cancer types (Fig. 4f). In order to account for cancer type confounding, we performed multivariate linear regression and showed that low-hypodiploids have significantly higher CNH and segment counts than both diploid and less-extreme aneuploid tumours after controlling for cancer type (*p <* 1e-17). This reinforces the association between high levels of chromosomal instability and hypodiploidy both between and within cancer types.

Interestingly, near-haploid tumours displayed significantly fewer intrachromosomal copy number alterations (*p <* 0.01, Fig. 4c) and lower intra-tumour copy number heterogeneity (*p <* 0.001, Fig. 4e) than low-hypodiploids, despite having higher rates of WGD. They also showed reduced frequency of chromothripsis (33% vs 55%, *p* = 0.03, Fig. S3e).

### Stable Hypodiploidy in Kidney and Adrenocortical Tumours Diverges from Pan-Cancer Patterns

The degree of chromosomal instability in hypodiploid tumours differs markedly between tissues. We previously identified two TCGA cancer types (KICH and ACC) with very high rates of hypodiploidy (Fig. 1b) and stereotyped chromosome loss patterns (Fig. 3b, Fig. S2c). Low-hypodiploids from these cancer types are further distinguished by having significantly lower intra-tumour copy number heterogeneity than low-hypodiploids from cancer types with non-stereotyped chromosome loss patterns (median 0.036 vs 0.103, *p <* 2.2 × 10^−16^, Fig. 4g). Moreover, they have significantly fewer intrachromosomal copy number alterations (median 107 vs 194 copy number segments per genome, *p <* 2.2 × 10^−16^, Fig. S3g), and the longest contiguous copy number segment of their average autosome spans 68% of the chromosome, compared to 48.2% in non-stereotyped hypodiploids.

The MH score heuristic, devised for ALL, is sensitive to karyotypic stability after genome doubling. We used this feature to assess chromosomal instability in hypodiploid tumours that have undergone WGD. We found that the heuristic detects 94.7% of doubled KICH hypodiploids and 87.5% of doubled ACC hypodiploids up to the near-triploid range (≤ 76 autosomes), versus only 19.7% of hypodiploids from non-stereotyped hypodiploid cancer types (Fig. S3h). These results indicate that hypodiploid ACC and KICH tumours are markedly more stable than hypodiploid cases from non-stereotyped TCGA tumours, even within the subset of cases that have undergone WGD, with stability comparable to hypodiploid ALL.

### Hypodiploidy is Associated with Poor Prognosis Across Cancer Types

Although hypodiploidy is an established predictor of poor prognosis in ALL and certain other cancer types including multiple myeloma^33^ and Hürthle cell carcinoma,^18^ its implications for survival in other cancer types are less clear. We assessed overall survival in relation to ploidy status across cancer types and found that survival was highest among patients with diploid tumours, followed by those with less-extreme aneuploid tumours (Fig. 4h). The overall survival of patients with low-hypodiploid tumours tracks closely with that of patients with polyploid tumours, in line with the known relationship between high copy number heterogeneity and poor prognosis.^7^

In a multivariate Cox proportional hazards regression controlling for age, sex, race, and cancer type, low-hypodiploidy was significantly associated with worse overall survival than both diploid and less-extreme aneuploid tumours (*p* = 0.024, *p* = 0.018 respectively). Although near-haploid tumours showed worse prognosis than diploid tumours when controlling for age, sex and race, this association was no longer significant when cancer type was included as a covariate (*p* = n.s.), although this may be due to limited sample size (N = 27).

We found no difference in overall survival between genome-doubled and non-doubled low-hypodiploid cases (*p* = 0.588, Fig. 4i), which supports the hypothesis that WGD is enriched in low-hypodiploid tumours due to their underlying chromosomal instability rather than a dosage effect increasing the fitness of the tumour.

### Hypodiploid Tumours are Enriched for TP53 Mutations and Hypoxia

Finally, we sought to shed light on the sources of the genomic instability observed in hypodiploid cancer genomes by analysing the genomic and environmental correlates of hypodiploidy. Overall, low-hypodiploid tumours have higher median mutation counts and ploidy-corrected mutation rates than diploid tumours, but are less likely to be hypermutated (Fig. S4a). Near-haploids do not differ significantly from diploids in mutation count.

We performed multivariate linear regression to detect associations between ploidy and mutations in genes from the COSMIC Cancer Gene Census,^34^ controlling for total mutation burden and cancer type (Fig. 5a). Low-hypodiploid tumours were strongly enriched for TP53 mutations compared to diploid tumours (log-odds 2.82, p_adj_ = 1.61e-46). Non-synonymous *TP53* mutations occurred in 69.4% of low-hypodiploid cases across the cohort, compared to 54.2% of near-haploids, 52.4% of polyploids, 31.5% of less-extreme aneuploids and only 11.6% of diploid cases (Fig. 5b). Low-hypodiploid tumours were also significantly depleted for mutations in *ARID1A, CTCF, ACVR2A, RPL22* and *BRAF* (p_adj_ *<* 0.05) compared to diploid tumours. All six of the genes differentially mutated in low-hypodiploid tumours showed similar patterns in polyploid tumours, where they were also significantly enriched or depleted, respectively (*p*_*adj*_ *<* 0.05). Moreover, across all COSMIC cancer genes, there was a weak positive correlation between the log odds ratios of gene mutation in hypodiploids vs polyploids overall (Fig. 5c). These results suggest that mutations in these genes affect generalised chromosomal instability or tolerance to aneuploidy rather than WGD specifically. There was no significant difference between low-hypodiploids and less-extreme aneuploid cases in the mutation rate of *ARID1A, CTCF, ACVR2A, RPL22* or *BRAF*, but low-hypodiploids remain enriched for *TP53* mutations even against other aneuploid cases (estimate 1.74, p_adj_ = 3.38 × 10^−41^). No genes were significantly differentially mutated in near-haploids compared to low-hypodiploids after multiple testing correction.

**Figure 5:**
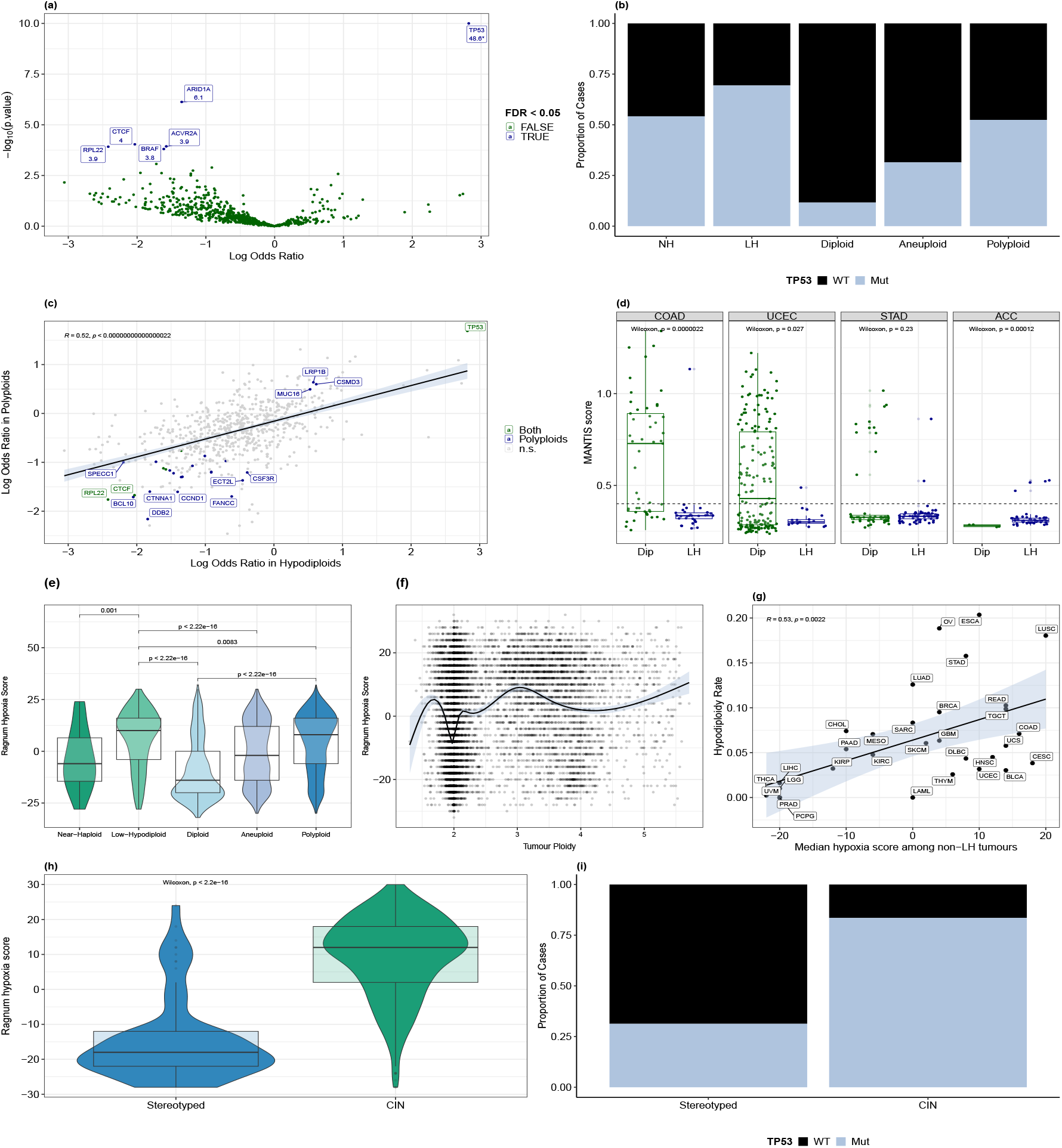
Hypodiploidy is associated with TP53 mutations and hypoxia. **A**, Genes enriched or depleted for mutations in low-hypodiploid vs near-diploid tumours, based on logistic regression controlling for total non-synonymous mutation count and cancer type. −log_10_(p-values) for genes significant after Benjamini-Hochberg correction are indicated below gene name labels. TP53’s −log_10_(p-value) is 63.5 but has been capped at 10 for visibility. A group of outliers with estimate ≤ −10 and p *>* 0.99 have been removed. **B**, Prevalence of non-synonymous TP53 mutations by ploidy class. **C**, Comparison between regression estimates (log odds ratio) for genes enriched/depleted in low-hypodiploid and polyploid tumours vs near-diploids. **D**, Distribution of microsatellite instability as measured by MANTIS score in colon (COAD), endometrial (UCEC), stomach (STAD) and adrenocortical (ACC) cancers. Dip, near-diploid; LH, low-hypodiploid. Dashed line indicates threshold for MSI-H status (MANTIS *>* 0.4). Points are jittered horizontally for visibility. **E**, Distribution of Ragnum hypoxia score by ploidy class. **F**, Continuous relationship between ploidy and Ragnum hypoxia score. **G**, Correlation between median Ragnum hypoxia score in non-hypodiploid tumours and rate of hypodiploidy across TCGA cancer types. **H**, Distribution of Ragnum hypoxia score in low-hypodiploid tumours from cancer types with stereotyped (KICH, ACC) vs non-stereotyped (‘CIN-driven’) hypodiploidy. **I**, TP53 non-synonymous mutation rate in low-hypodiploids from cancer types with stereotyped vs non-stereotyped hypodiploidy. All analyses in this section were performed on the TCGA dataset.

While RPL22 mutations can impair TP53 function,^35^ mutations in this gene are also associated with microsatellite instability (MSI), which is known to be mutually exclusive with genome doubling in cancer.^6,5,20^ We extended this observation to low-hypodiploid tumours by analysing microsatellite instability using MANTIS scores from Bonneville et al (2017).^36^ In all three of the canonical MSI cancer types (colon, endometrial, stomach), low-hypodiploid tumours were less likely to display a high MSI phenotype (MANTIS *>* 0.4) (Fig. S4c), and in colon and endometrial cancer, LH tumours had significantly lower MANTIS scores overall (Fig. 5d). In adrenocortical carcinoma, however, which has both an appreciable rate of microsatellite instability and stereotyped hypodiploidy, hypodiploid tumours have higher MANTIS scores and are more likely to be classified as microsatellite-unstable (Fig. 5d, Fig. S4c). The same pattern is observed even when genome-doubled low-hypodiploid tumours are excluded, suggesting that it is not merely a result of the known anticorrelation between microsatellite instability and WGD (Fig. S4d).

Tissue hypoxia has been linked to replication stress, DNA repair deficiency and chromothripsis.^37^ To investigate the relationship between hypoxia and large-scale aneuploidy, we computed hypoxia scores using the Ragnum pimonidazole^38^ gene signature across all TCGA samples. Low-hypodiploid tumours were the most hypoxic of any ploidy class, followed by polyploid tumours (Fig. 5e). Interestingly, hypoxia scores varied continuously with ploidy, peaking around ploidy values of 1.5 and 3 (triploidy). Perfectly-diploid tumours were the least hypoxic, with perfectly-tetraploid tumours also having relatively low hypoxia scores (Fig. 5f). To account for the possibility that highly-aneuploid tumours express higher levels of hypoxia-related genes for other reasons, we calculated the median hypoxia score per cancer type and compared it to the rate of hypodiploidy, finding a strong positive correlation (r = 0.53, *p* = 0.0022, Fig. 5g). There were significant positive correlations between median hypoxia score and both hypodiploidy and polyploidy across 5 of 8 hypoxia signatures computed by Bhandari et al. (2019)^39^ (Fig. S4g).

We considered the possibility that this relationship is confounded by replication rate, as a high tissue replication rate is associated with polyploidy across cancer types and fast-growing tumours are more likely to become hypoxic. While low-hypodiploidy rate is correlated with median proliferative index across tissues (r = 0.36, *p* = 0.044, Fig. S4h), linear regression of hypoxia score on ploidy class reveals that hypodiploid tumours are still significantly more hypoxic than both diploid and aneuploid tumours after taking proliferative index into account (*p* = 1.63e-35 and 0.0127, respectively).

Notably, the two TCGA cancer types with stereotyped low-hypodiploid chromosome loss patterns were an exception. Low-hypodiploid cases of kidney chromophobe and adrenocortical cancer had much lower hypoxia than low-hypodiploids from other cancer types (Fig. 5h), and these cancer types had the lowest and eighth-lowest median tissue hypoxia levels of all 33 cancer types (Fig. S4e). Additionally, low-hypodiploid cases of the stereotyped hypodiploid cancers had lower *TP53* mutation rates: *TP53* is non-synonymously mutated in 23.4% of low-hypodiploid ACC and 38.5% of low-hypodiploid KICH, versus a median of 85.7% in low-hypodiploids from other cancer types (Fig. 5i).

## DISCUSSION

In recent years, much work has focused on the antecedents, correlates and outcomes of extreme aneuploidy in cancer, particularly via genome doubling (e.g.^5,20, 6,40, 41^). However, outside specific cancer types with stereotyped hypodiploid subtypes (e.g.^8,18, 13, 42^), the occurrence of extreme aneuploidy and genomic imbalance through chromosome loss has received relatively little attention. Here, we have identified hypodiploidy as a widespread phenomenon across cancer types and systematically characterised patterns of chromosome loss and genomic instability in a pan-cancer framework.

### A Generalised Chromosomal Instability Phenotype

Beyond their defining characteristic—high levels of whole-chromosome aneuploidy—low-hypodiploid tumours exhibit dramatically increased chromosomal instability at multiple scales, from smallscale intrachromosomal copy number alterations and chromothripsis to whole genome duplication. These tumours also have very high levels of intratumour copy number heterogeneity, suggesting ongoing instability rather than stable maintenance of defined hypodiploid karyotypes. While the hypodiploid state may promote other forms of chromosomal instability, it has been shown that monosomies alone do not induce CIN,^43^ and cross-tissue correlations suggest that these features are manifestations of an underlying CIN phenotype.

Like genome-doubled tumours,^6^ low-hypodiploids are strongly enriched for *TP53* mutations^8^ and depleted for mutations in microsatellite instability-related genes. Low-hypodiploids are also significantly more hypoxic than diploid and less-extreme aneuploid tumours, and rates of both low-hypodiploidy and polyploidy across cancer types correlate with median tissue hypoxia levels. Hypoxia induces replication stress and suppresses DNA damage/repair responses;^44^ it has been linked to centrosome aberrations,^45^ homologous repair deficiency, loss of heterozygosity,^46^ increases in mutational burden, chromothripsis and aneuploidy.^39, 47^ The commonalities between factors associated with low-hypodiploidy and polyploidy suggest that these factors do not specifically cause genome doubling but instead promote generalised chromosomal instability and/or enable tolerance to extreme aneuploidy.

Intriguingly, near-haploid tumours do not show the same CIN patterns as low-hypodiploids. While our sample size for near-haploids was limited, we observed significantly fewer intrachromosomal copy number events and lower levels of chromothripsis and intra-tumour copy number heterogeneity than low-hypodiploid tumours. Near-haploid tumours were also less hypoxic than low-hypodiploids, and the prevalence of near-haploidy appears generally unrelated to that of low-hypodiploidy across cancer types. These data support the hypothesis that near-haploidy arises through a distinct mech-anism, such as direct near-haploidisation,^48^ while low-hypodiploids typically result from persistent chromosome missegregation and are thus enriched in cancer types that tend to be chromosomally unstable.

In contrast to the general trend of extreme chromosomal instability among low-hypodiploid tumours, we identified a subset of cancer types with stereotyped low-hypodiploid chromosome loss patterns: ALL, kidney chromophobe, and adrenocortical carcinoma. KICH and ACC were notable for having very high rates of low-hypodiploidy, at 80% and 54.4% compared to 20.3% in the next-highest cancer type, oesophageal carcinoma. Hypodiploidy is an established phenomenon in KICH, with stereotyped loss of chromosomes 1, 2, 6, 10, 13, 17 and, less commonly, 21.^12^ Hypodiploidy was also identified as one of three copy number subtypes in the TCGAACC cohort, with frequent genome doubling and preservation of LOH patterns in genome-doubled cases.^19^

These stereotyped hypodiploid tumours show significantly reduced intra-chromosomal instability, intra-tumour copy number heterogeneity, hypoxia, *TP53* mutation burden and post-WGD missegregation compared to non-stereotyped low-hypodiploids. Hyperdiploid ALL has previously been established as an example of stable aneuploidy;^30^ this research supports the existence of (relatively) stable extreme aneuploidy after chromosome loss as well. We thus distinguish two classes of hypodiploid tumour: stereotyped hypodiploids, in which hypodiploidy is stable and likely selected for, and sporadic hypodiploids, which represent the extreme end of a spectrum of chromosomal instability. Our analysis suggests that even extreme levels of chromosomal instability rarely result in near-haploid karyotypes or in hypodiploidy rates above ~20 %.

### The MH Score and the Diagnostic Challenge of Masked Hypodiploid ALL

Genome doubling presents an important diagnostic challenge in hypodiploid ALL, because it can obscure poor-prognosis masked hypodiploids with apparent good-prognosis high-hyperdiploid karyotypes, risking treatment failure.^11^ Previous research has addressed this problem using loss patterns of specific chromosomes^49^ or targeted FISH/flow-cytometry,^50, 51^ but these approaches either miss near-haploid cases or rely on the persistence of a hypodiploid subclone. Additionally, approaches based on the relative copy numbers of specific chromosomes are likely to suffer from the fact that chromosomes retained in hypodiploids also tend to be gained in hyperdiploids.^30^

We thus developed and tested a simple heuristic, the MH score, to distinguish masked hypodiploid from high-hyperdiploid ALL using only cytogenetic information, even in cases of complete masking. The MH score is based on the observation that masked hypodiploids tend to exhibit ‘hyperdiploidy by tetrasomy’, with genomes composed mostly of disomies and tetrasomies, but not trisomies like in hyperdiploid ALL.^22, 11^ This pattern is expected following a WGD event on a hypodiploid background. However, chromosomal instability post-doubling can degrade the pattern over time, as missegregation events create additional aneuploidies, including trisomies. Indeed, in more chromosomally unstable tumours, the MH score heuristic loses discriminative power, and masked hypodiploids may be indistinguishable from hyperdiploid/near-triploid cases. It was thus important to test the performance of this heuristic in ALL, and to characterise its sensitivity and specificity in detail across the full range of doubled hypodiploid chromosome counts.

Using a combination of allele-specific copy number data and multi-clone phylogenetic trees, we showed that the MH score is highly specific (*>*99%) in the clinically relevant high-hyperdiploid autosome count range and sensitive (~90%) across the range of potential doubled hypodiploid autosome counts. Sensitivity falls to 82.5% in the high-hyperdiploid range, and the score heuristic shows low specificity for cases with 46 autosomes, where few missegregation events are required to shift the tetrasomy/trisomy balance. The false negative cases may reflect a chromosomally-unstable subset of ALL or older WGD events. These cases were enriched for low-hypodiploids, supporting the idea that low-hypodiploids are more chromosomally unstable than near-haploids even in ALL. We conclude that the MH score should be a useful tool for analyses of fully-masked hypodiploid ALL using only cytogenetic data. However, we caution that the absence of a ‘hyperdiploidy by tetrasomy’ pattern does not definitively preclude a hypodiploid history, particularly in genomically-unstable cases, and recommend increased adoption of allele-specific genotyping in clinical diagnostic contexts where feasible.

### The Fitness of Hypodiploid Tumours

Across the TCGA cohort, tumour hypodiploidy is associated with poor prognosis, suggesting increased tumour aggressiveness. It is not clear whether this is due to their absolute ploidy or to their chromosomal instability, but we observed that the survival curve for low-hypodiploid tumours overlaps that of polyploid tumours, which have opposite ploidy but similar levels of chromosomal instability. Previous research has shown that intratumour copy number heterogeneity is a stronger prognostic factor pan-cancer than the mere presence of copy number alterations,^7^ and chromosomal instability correlates with disease progression across aneuploid paediatric B-ALL xenografts.^52^ The poor prognosis of hypodiploid tumours may therefore be attributable to their underlying genomic instability providing a rich substrate for selection.^53^

It is plausible that extensive chromosome losses and the resulting genomic imbalances alter signalling networks,^54^ create a sensitised background for tumour suppressor gene mutations,^18^ or induce metabolic changes.^55^ Analysis of chromosome loss patterns in low-hypodiploid tumours reveals no evidence of selection to avoid loss of large or dosage-sensitive chromosomes. The only factor that consistently predicted chromosome loss rate was driver gene density. This finding is consistent with a model in which driver aneuploidies occur early, potentially before the establishment of chromosomal instability, and are followed by unconstrained chromosome losses without regard to the level of genomic imbalance created. Given the inefficiency of negative selection in cancer, particularly in cases with high mutational burdens, it remains unclear to what extent this level of aneuploidy imposes a fitness cost on the tumour.^56^

It has been proposed that genome doubling is selected in hypodiploids and in tumours with widespread LOH in order to restore gene dosage and buffer deleterious mutations.^11, 20^ However, we observed no difference in tumour aggressiveness between doubled and non-doubled low-hypodiploids, and suggest that the increased WGD rate in hypodiploid tumours may reflect a generalised chromosomal instability phenotype that drives both chromosome loss and genome doubling. We were not able to assess the relationship between tumour fitness and near-haploidy directly, but it is plausible that genome doubling is more beneficial to these cases. Supporting this, we find that genome doubling is strongly enriched in near-haploid tumours despite their significantly lower levels of other forms of chromosomal instability. However, our MH score analyses suggest that the doubled clone fully outcompetes the hypodiploid clone in only a minority (≲ 15%) of apparently-hyperdiploid ALL cases in the Mitelman database. Importantly, near-haploid clones can persist throughout tumour evolution and regularly give rise to highly-aggressive tumours and relapses.

Our pan-cancer analysis reveals that most hypodiploid tumours are characterised by extreme chromosomal instability at multiple scales, distinguishes a subset of stable hypodiploids with characteristic chromosome loss patterns, and provides an illustrative example of the extreme freedom from dosage constraint under which tumours evolve.

**Figure S1:**
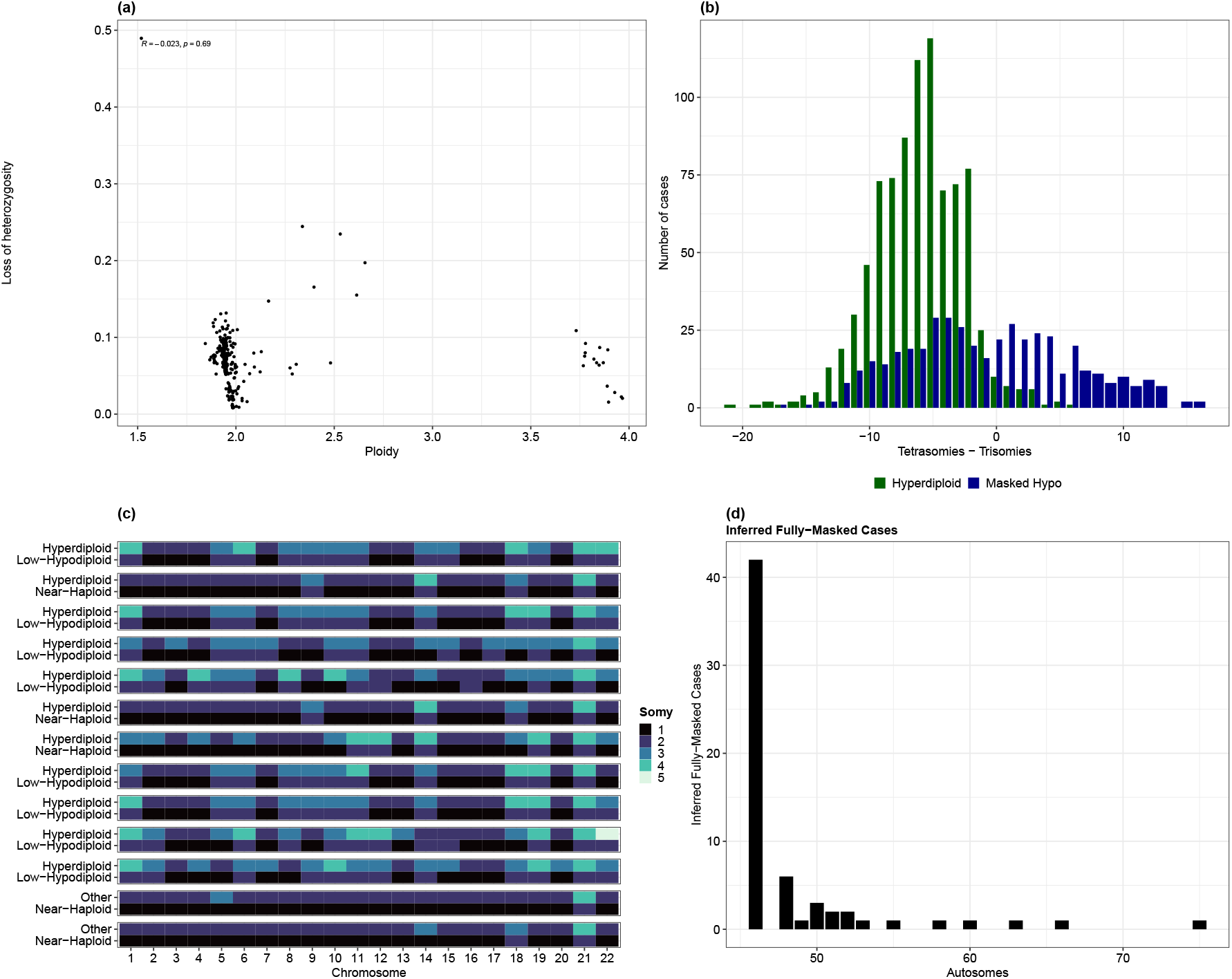
Related to Fig. 2. Identification of masked hypodiploidy based on allele-nonspecific data. **A**, Distribution of ploidy and loss of heterozygosity among 293 cases from the TARGET-ALL-P2 project. Hypodiploid cases with multiple WGD events would appear in the top right. **B**, Distribution of MH score (# tetrasomies - # trisomies, excluding the sex chromosomes) in the TCGA cases. Masked hypodiploid cases include any near-haploid or low-hypodiploid TCGA case with a positive WGD call; hyperdiploid cases were defined based on the absence of a hypodiploid history and an autosome count between 49 and 65 (adjusted from 51-67 chromosomes in Woodward et al). **C**, Visualisation of chromosome counts in false negative cases. Each facet represents a case with two clones; colour indicates chromosome copy number (somy). **E**, Autosome counts of inferred fully-masked hypodiploids in ALL from the Mitelman database.

**Figure S2:**
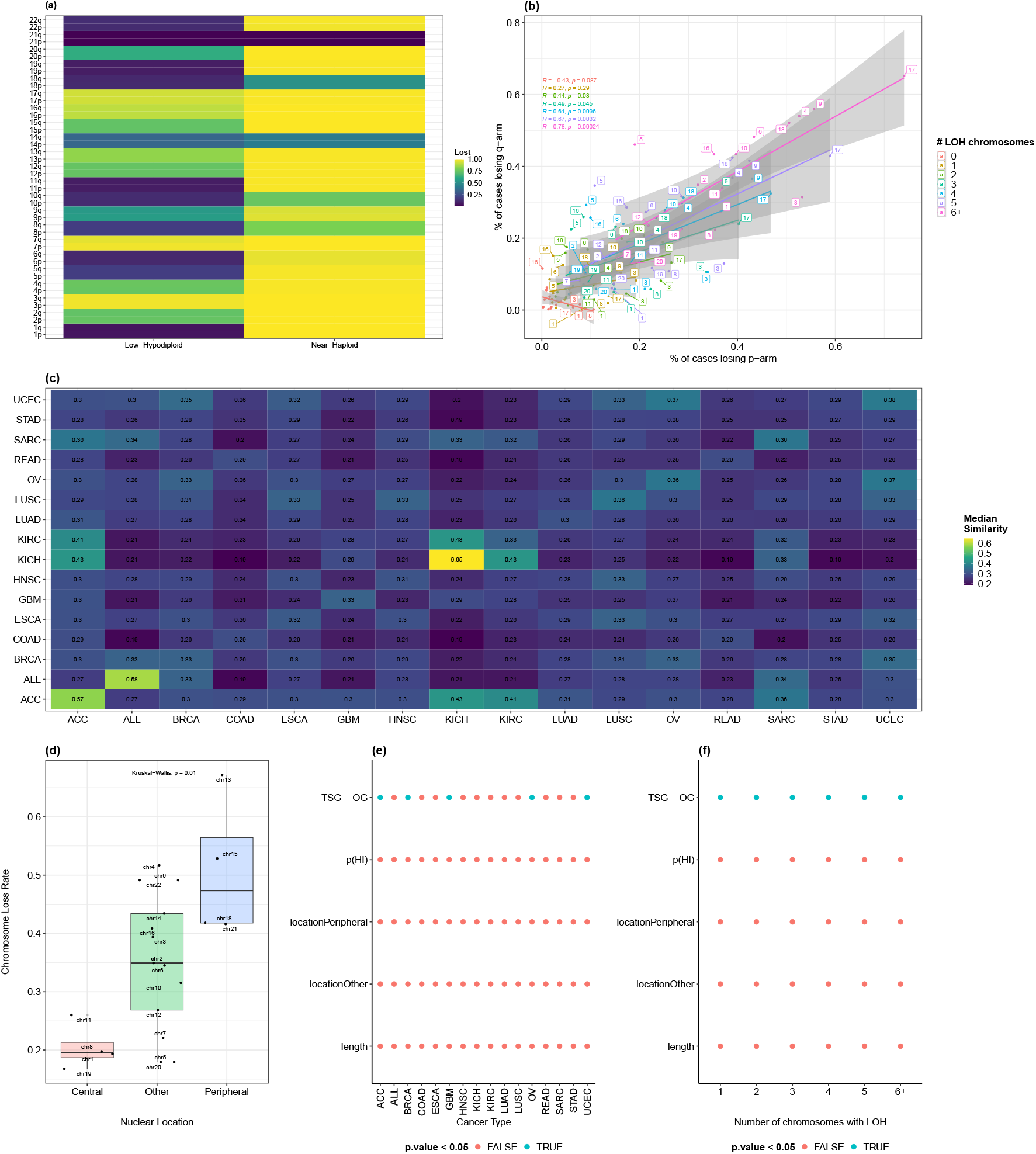
Related to Fig. 3. Patterns of chromosome loss in hypodiploid tumours. **A**, Chromosome arm loss rates in near-haploid and low-hypodiploid ALL from the Mitelman database; colour indicates the proportion of cases in which a given chromosome is in one copy. **B**, Correlation between loss rates of p and q chromosome arms within aneuploid tumours in the TCGA dataset, separated by the number of whole chromosomes with LOH–6+ represents hypodiploid cases. **C**, Median pairwise Jaccard similarity between samples of different cancer types. Intra-tissue similarities are shown on the diagonal. **D**, Pan-cancer chromosome loss rate in low-hypodiploid tumours by nuclear location, excluding chromosome 17. **E**, Tissue-specific generalised linear regression of loss rate in low-hypodiploids on chromosome features including length in Mb, nuclear location (with ‘central’ as reference) from Parl et al (2023), TSG - OG density score from Davoli et al, and dosage sensitivity, measured as the sum of probabilities of haploinsufficiency (pHI) from Collins et al (2022) for each gene on the chromosome. **F**, Generalised linear regression of loss rate on chromosome features, split by number of chromosomes with LOH (1, 2, 3, 4, 5 and 6+). ALL was excluded from (f) due to the lack of LOH information in the Mitelman database.

**Figure S3:**
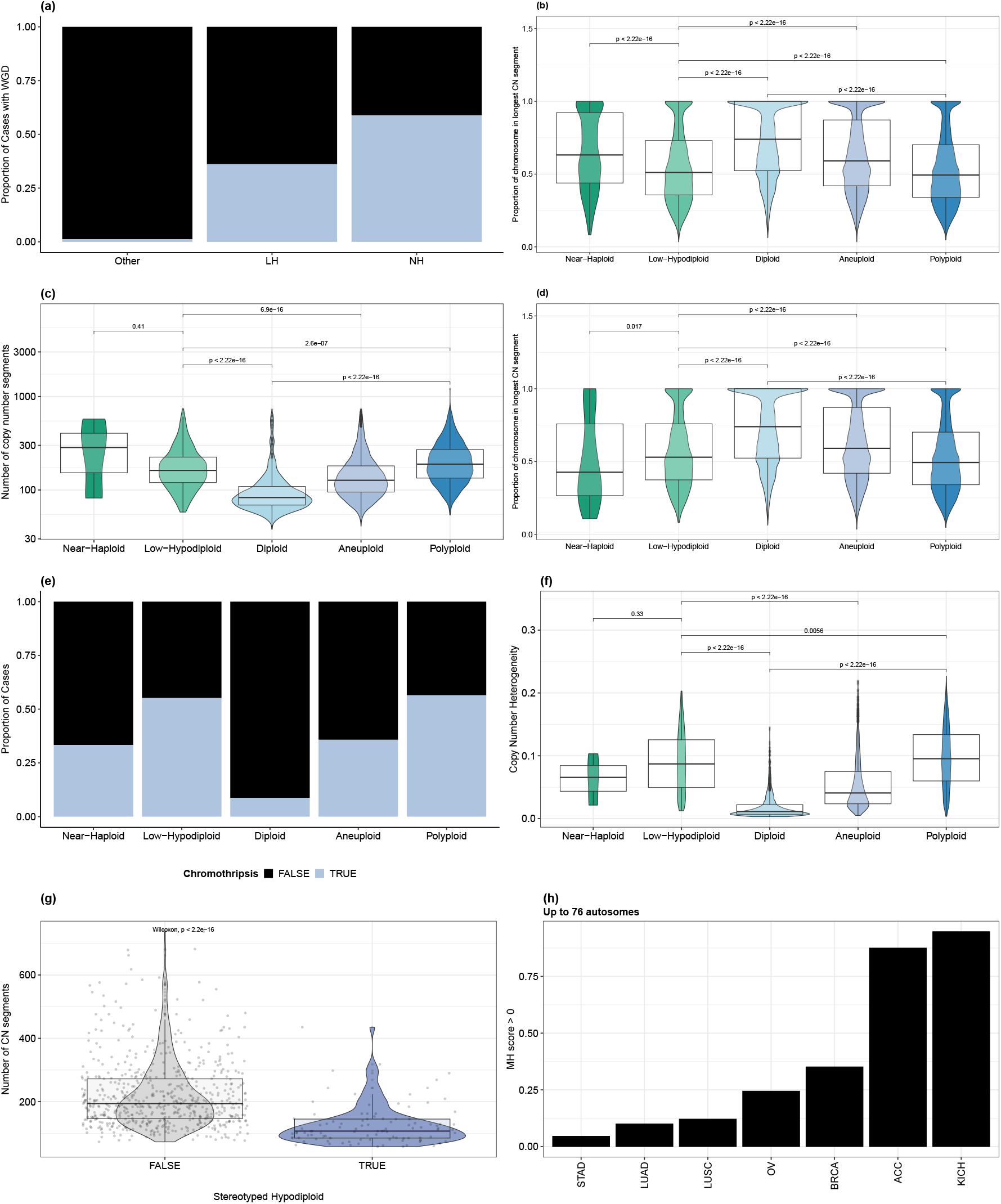
Related to Fig. 4. Hypodiploid tumours are distinguished by chromosomal instability at multiple scales. **A**, WGD rate by hypodiploidy level in the Mitelman ALL dataset, but clones with MH score *>* 0 and autosome count outside the hypodiploid range were reassigned to the ‘Other’ group and not called as WGD unless they met the ploidy criterion. Multi-clone cases were assigned their lowest ploidy class. **B**, Proportion of chromosome in its longest contiguous segment by class. **C**, Number of copy number segments (log10-transformed) by class. **D**, Proportion of chromosome in its longest contiguous segment by class, excluding doubled hypodiploids. **E**, Proportions of TCGA cases assigned as chromothriptic by Rasnic & Linial (2021) by ploidy class. **F**, Distribution of copy number heterogeneity by ploidy class, excluding genome-doubled hypodiploids *(cont*.) **G**, Distribution of segment count in low-hypodiploid cases from cancer types with stereotyped and non-stereotyped chromosome loss patterns (KICH and ACC vs BRCA, LUSC, OV, STAD, LUAD, ESCA, COAD, HNSC, KIRC, SARC, UCEC, READ, GBM). **H**, Sensitivity of the MH score heuristic (tetrasomies - trisomies *>* 0) in classifying TCGA samples with a hypodiploid history, a positive WGD call and ≤ 76 autosomes as masked hypodiploids. Only cancer types with ≥ 15 doubled hypodiploids with ≤ 76 chromosomes are included.

**Figure S4:**
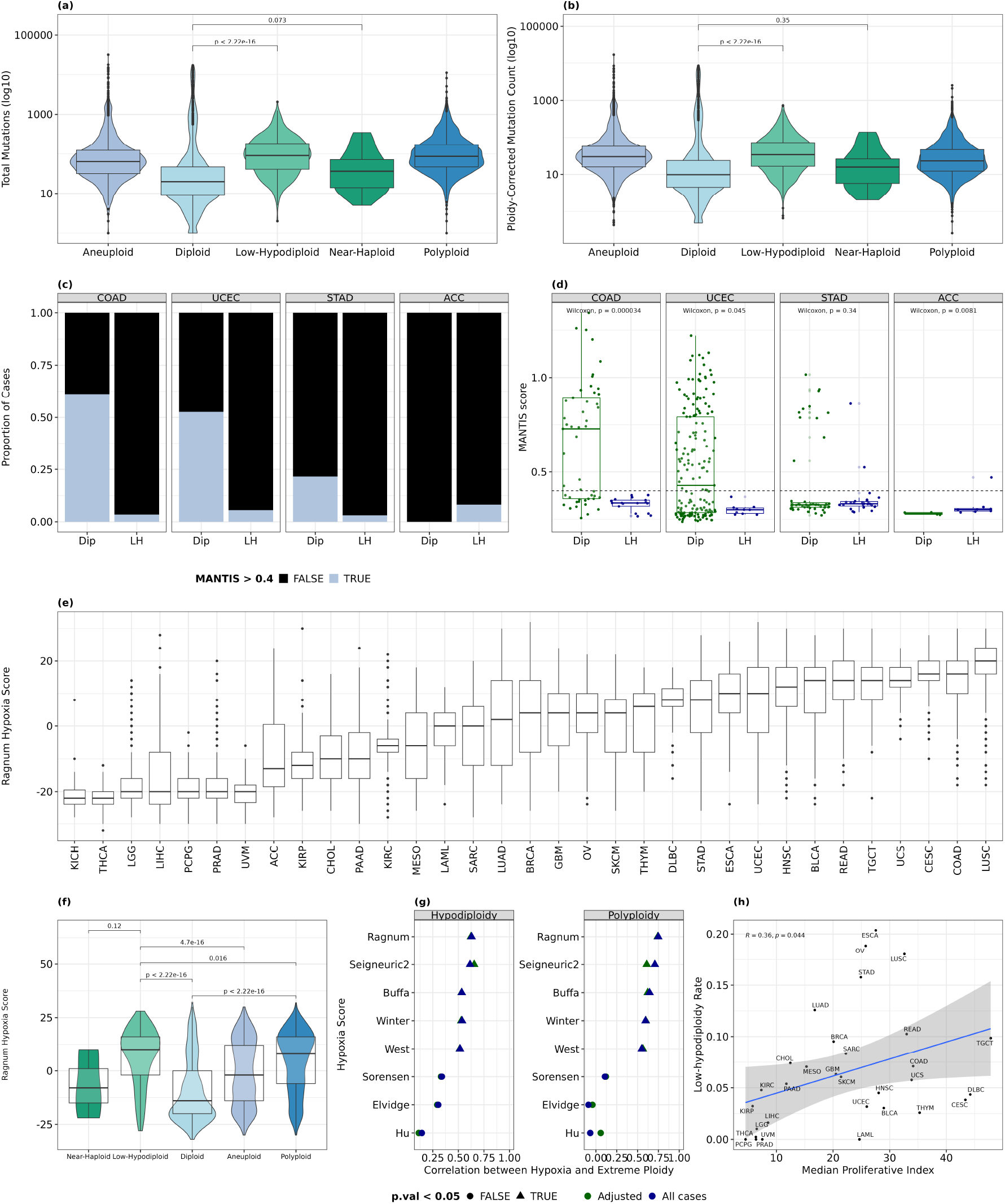
Related to Fig. 5. Hypodiploidy is associated with TP53 mutations and hypoxia. **A**, Total mutation counts by ploidy class. **B**, Mutation rates by ploidy class, mutation counts divided by ploidy. **C**, Proportion of cases with MSI-H status (microsatellite instability - high), defined by a MANTIS score *>* 0.4, in colon (COAD), endometrial (UCEC), stomach (STAD) and adrenocortical (ACC) cancers. Dip, near-diploid; LH, low-hypodiploid. **D**, Distribution of MANTIS scores after removing genome-doubled hypodiploid tumours. Points are jittered horizontally for visibility. **E**, Distribution of Ragnum hypoxia scores by cancer type, including all ploidy classes. **F**, Ragnum hypoxia score by ploidy class after removing genome-doubled hypodiploid tumours. *(cont*.) **G**, Correlations between median hypoxia score and rate of hypodiploidy or polyploidy based on eight different hypoxia signatures. Adjusted points indicate correlations based on median hypoxia scores calculated using non-hypodiploid/non-polyploid cases respectively.**H**, Cross-tissue correlation between low-hypodiploidy rate and median proliferative index. ACC and KICH were excluded due to their outlying hypodiploidy rates. Proliferative index was calculated based on the median CPM-normalised expression of genes in the metaPCNA proliferation signature from Venet et al (2011).

## METHODS

### Copy Number Data

We downloaded all acute lymphoblastic leukaemia karyotypes from the Mitelman Database of Chromosome Aberrations & Gene Fusions in Cancer and used CytoConverter,^26^ bedops partition and bedtools map to calculate net copy number alterations for each genomic region. We removed cases with errors from CytoConverter or with unknown chromo-some numbers, then calculated copy number as 2 + net copy number alteration. Cases with 46 chromosomes and no numerical abnormalities were inserted separately as diploids. For the TCGA tumours, allele-specific copy number profiles computed using ASCAT were downloaded using TCGAbiolinks; we removed metastatic and recurrent tumours, FFPE and ASCAT-defined suspected contaminated samples, and deduplicated to keep one sample per patient. We ran MEDICC2 on ASCAT allele-specific absolute copy number profiles for all TCGA samples and defined a sample as WGD-positive if the minimum event distance was decreased when WGD was included as a possible evolutionary step. Chromo-some somy was calculated as the mean copy number along the chromosome, rounded to a whole number. Samples with nullisomic chromosomes or sub-haploid autosome counts were excluded. Autosome counts were used instead of chromosome counts due to the unreliability of sex chromosome copy number assignments in these datasets.

### Identifying Hypodiploids

The threshold for hypodiploidy was set at ≥ 6 autosome losses, which corresponds to ≤ 38 autosomes. This threshold was chosen based on the distributions of autosome counts in two cancer types with established hypodiploid subtypes (ALL and KICH). Samples with 23-27 autosomes were classified as near-haploid, while samples with 28-38 autosomes were classified as low-hypodiploid. Tumours with 39-43 autosomes were considered high-hypodiploid and grouped with the less-extreme aneuploid samples in this analysis. Samples with 49-65 autosomes were classified as hyperdiploid based on the 51-67 chromosome range defined for ALL,^30^ subtracting two due to the lack of sex chromosomes.

In the TCGA dataset, chromosome losses were as-certained using LOH information, which allowed us to identify former hypodiploids that had since undergone WGD or otherwise gained chromosomes. Autosomes with LOH over *>*90% of their length were defined as ‘lost’ and assigned a somy_min_ of 1, while other autosomes were assigned a somy_min_ of 2. Minimal autosome count (autosome_min_) during the tumour’s evolution was computed as the sum of somy_min_ across all autosomes. While individual LOH events can occur after WGD if the same homolog is lost twice, meaning that the tumour was never necessarily in a hypodiploid state, we assume that LOH events occurred pre-WGD and count tumours with an autosome_min_ ≤ 38 as (former) hypodiploids. Where LOH information was unavailable, i.e. in the Mitelman-ALL dataset, hypodiploids were identified based on current autosome count at time of karyotyping, calculated as the sum of rounded chromosome ploidies.

### Distinguishing Masked Hypodiploids from Hyperdiploids using Cytogenetic Data

We calculated a masked hypodiploidy (MH) score for each sample as # tetrasomic chromosomes - # trisomic chromosomes. To test the ability of the score to distinguish masked hypodiploids from hyperdiploids, we gathered two datasets of masked hypodiploids (positives) and hyperdiploids (negatives). For the positives, we took Mitelman ALL cases that had both a hypodiploid and a non-hypodiploid clone, built copy number phylogenetic trees with MEDICC2 and selected cases where WGD was inferred and the non-hypodiploid clone had at most 76 autosomes (N = 123). For the negatives, we obtained allele-specific hyperdiploid copy number data from Woodward et al (2023) (N = 577) and counted chromosomes labelled as “Tri” vs “Tetra 2 2” or “Tetra 3 1” after excluding the sex chromosomes and chromosomes with subclonal copy number variation. We additionally used the TARGET-ALL-P2 dataset (N = 293, 292 non-hypodiploid) to assess accuracy outside the high-hyperdiploid range. ASCAT allele-specific copy numbers were downloaded, deduplicated per patient and hypodiploidy inferred as described for the TCGA dataset.

### Chromosome Loss Patterns

We created a binary loss/retention profile for each chromosome arm in each sample, defining a chromosome arm as lost if it had *>*90% LOH (TCGA samples) or had copy number *<* 2 for *>*90% of its length (Mitelman samples). The sex chromosomes and acrocentric arms (13p, 14p, 15p, 21p, 22p) were excluded due to lack of data. The average loss rate for each chromosome arm was calculated separately by tissue. We clustered all samples by their chromosome loss profiles using the Jaccard distance (1 - number of shared lost chromosome arms divided by total number of lost chromosome arms between the two samples), and calculated the median Jaccard similarity (1 - D) within and between each cancer type. The heatmap was created with ComplexHeatmap in R. To investigate factors associated with chromosome loss rate, we obtained chromosome tumour suppressor - oncogene density scores from Davoli et al (2013), chromosome nuclear locations inferred based on interchromosomal translocation frequency by Parl et al (2023), chromosome lengths from the D3GB R package, and gene haploinsufficiency probabilities from Collins et al (2022).^57^ These haploinsufficiency probabilities were summed for each chromosome to create a chromosomal dosage sensitivity score. We performed linear regression of chromosome loss rate on chromosome length, nuclear location, driver gene density and dosage sensitivity score pan-cancer and separately by tissue using glm() in R. To assess whether the overall level of aneuploidy affects these relationships, we re-ran the pan-cancer regression analysis for the TCGA tumours after stratifying samples by their number of chromosome losses (1, 2, 3, 4, 5, 6+, where 6+ indicates current or former hypodiploid samples). Cancer types were defined as stereotyped hypodiploids if the median Jaccard similarity between low-hypodiploid chromosome arm loss profiles from tumours of that cancer type was higher than all median similarities between that cancer type and other cancer types. For the ALL cases, only current hypodiploid cases (autosome count ≤ 38) were included.

### Chromosomal Instability

TCGA cases were defined as WGD-positive if they had a positive MEDICC2 WGD call. ALL cases were defined as WGD-positive if they had ploidy *>* 2.7, if they had a hypodiploid and non-hypodiploid clone and a positive MEDICC2 WGD call, or if they had an MH score *>* 0. Fully-masked clones (MH score *>* 0, no current-hypodiploid subclone) were classified as former near-haploids or former low-hypodiploids based on their autosome count divided by two. For the MH score criterion, we initially included all inferred fully-masked clones; in a supplementary analysis, clones with MH score *>* 0 but autosome count outside the well-characterised hyperdiploid range were excluded as masked hypodiploids and reclassified as non-hypodiploid and non-doubled (unless eligible under the ploidy criterion). Cases with any WGD-positive clones were counted as WGD-positive.

Intrachromosomal instability was measured for each TCGA sample by counting the number of copy number segments in each sample, and by computing for each chromosome the proportion of the summed length of all segments occupied by its longest contiguous segment. Chromothripsis status for each TCGA sample was derived from Rasnic & Linial (2021)^58^ and copy number heterogeneity values from van Dijk et al (2021). When computing the correlation between WGD rate and hypodiploidy rate across cancer types, hypodiploid samples were excluded from the set of samples used to calculate WGD rate to disentangle the increased WGD rate in hypodiploid tumours from the hypothesised underlying chromosomal instability phenotype driving both WGD and hypodiploidy. Similarly, median segment counts and median copy number heterogeneity were calculated based on samples without gross ploidy alterations (hypodiploidy or polyploidy). Two cancer types, ACC and KICH, were excluded from these correlations due to their extremely high rates of hypodiploidy.

### Survival Analysis

Survival analysis was performed using Cox proportional hazards regression with the R survival and survminer packages as follows:

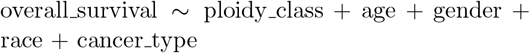

### Genomic Correlates of Hypodiploidy

We downloaded MAFs for each TCGA cancer type from the Genomic Data Portal using TCGAbiolinks and calculated the total number of mutations, the ploidy-corrected mutation rate (mutations / ploidy) and the number of non-synonymous mutations per sample. To detect genes enriched for mutations in low-hypodiploid samples compared to diploid samples, we performed logistic regression controlling for total non-synonymous mutation count and cancer type, using only genes present in the COSMIC Cancer Gene Census and with ≥ 5 total non-synonymously mutated samples across the low-hypodiploid and diploid cohorts. A similar analysis was repeated to compare polyploids to diploids, low-hypodiploids to aneuploids, and low-hypodiploids to near-haploid cases.

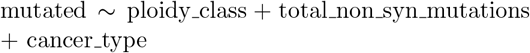

RNA-seq counts were downloaded from the Genomic Data Commons and normalised using cpm() in edgeR. We computed hypoxia scores for every TCGA sample using the Ragnum pimonidazole signature^38^ as described by:^39^ for each gene in the Ragnum signature, a sample was assigned a score of +1 if its expression for that gene was higher than the median expression of that gene across all samples and −1 otherwise, and scores were summed for each sample. To confirm that our results were robust to choice of hypoxia signature, we additionally analysed the relationship between hypodiploidy rate and hypoxia score using eight hypoxia scores (Ragnum, Buffa, Winter, West, Sorensen, Elvidge, Hu and Seigneuric2) precomputed by Bhandari et al (2019) for a subset of 20 cancer types. We computed correlations between hypoxia scores from each signature and both hypodiploidy and polyploidy. We also computed adjusted versions of these correlations by removing hypodiploid and polyploid samples (respectively) from the set of samples used to compute tissue median hypoxia scores. Microsatellite instability scores (MANTIS scores) were obtained from Bonneville et al (2017),^36^ and samples were classified as MSI-H if their MANTIS score was *>* 0.4. The proliferative index for each sample was calculated as the median expression (CPM) of 117 genes included in the metaPCNA proliferation signature. This signature was derived by Venet et al (2011)^59^ from the top 1% of genes correlated with the proliferation nuclear cell antigen across healthy samples.

### Statistical analysis

Multiple testing correction was performed by calculating the Benjamini-Hochberg False Discovery Rate; tests with FDR *<* 0.05 were considered significant. The analyses in this paper were performed using R (v4.2), data.table v1.16.2, tidyverse 2.0.0, janitor 2.2.0, TCGAbiolinks 2.34.0, TCGAutils 1.26.0, openxlsx 4.2.7.1, ggpubr 0.6.0, ggrepel

0.9.6, patchwork 1.3.0, ComplexHeatmap 2.22.0, ggplotify 0.1.2, RColorBrewer 1.1.3, survival 3.7.0, survminer 0.5.0, broom 1.0.7, DescTools 0.99.58, D3GB 2.0 and MicroViz 0.12.7.

## Acknowledgements

This research was funded by Research Ireland through the Research Ireland Centre for Research Training in Genomics Data Science under grant number 18/CRT/6214 (E.L.), by the European Research Council under grant agreement 771419 (A.McL), and by the HEA-HCI Pillar 3 initiative (M.N.L. and E.L.).

This project is based on data from The Cancer Genome Atlas and TARGET-ALL-P2 projects, as well as the Mitelman Database of Chromosome Aberrations and Gene Fusions in Cancer. The authors thank the patients who participated in these studies and their families. We also thank Tom Kaufmann for helpful discussions.

## Author Contributions

EL conceived of and designed the project, analysed and interpreted data, produced all main and supplemental figures and wrote the manuscript. MNL provided project direction, assisted with data interpretation, edited the manuscript, and provided funding. AMcL supervised the project and provided project direction and funding.

## Data and Code Availability

The scripts required to reproduce the analyses in this paper are available at https://github.com/loughrae/Hypodiploidy. This paper analyzes existing, publicly available datasets, whose accessions are also found in the key resources table.

